# *AMBER* and *GOLD*: Polycistronic Genes for Betaxanthin Production in Plants

**DOI:** 10.64898/2026.01.14.699277

**Authors:** Nicholas Desnoyer, Lionel Hill, Mark Youles, Sophien Kamoun

**Affiliations:** The Sainsbury Laboratory, University of East Anglia; Norwich Research Park, Norwich, UK; John Innes Centre, Norwich Research Park, Norwich, UK

## Abstract

Genetically encoded pigments are powerful visual reporters and creative tools for biology, yet in plants the palette of pigment biosynthesis genes has remained largely limited to red betacyanins encoded by *RUBY*. Here we develop and characterize three new polycistronic constructs *AMBER_v1, AMBER_v2*, and *GOLD* that contain betalain biosynthesis enzymes to produce yellow, fluorescent betaxanthins in plant tissues. These tools expand the palette of publicly available pigmentation genes for use in plant research, education, and floral design.

## INTRODUCTION

In plants, novel floral coloration, including blue chrysanthemums ^1^, orange petunias ^2^, and a broad range of fuchsia flowers ^3^ have been produced through genetic modification of pigment biosynthetic pathways. However, most have not been developed into practical reporter systems due to the complex genetic structure of their biosynthetic pathways. The most widespread gene used as a color reporter in plants is *RUBY*, a polycistronic construct encoding three beet (*Beta vulgaris*, Caryophyllales) genes of the betacyanin pathway, *Cytochrome P450 76AD1* (*CYP76AD1), L-DOPA, 4,5-dioxygenase* (*DODA*), and *Cyclo-DOPA 5-O-glucosyltransferase* (*GT)* separated by self-cleavable 2A linkers ^3^. Specifically, *CYP76AD1* hydroxylates tyrosine into L-DOPA and additionally oxidizes L-DOPA to cyclo-DOPA (Figure 1)^4,5^. L-DOPA is also converted to betalamic acid by L-DOPA, 4,5-dioxygenase (DODA) which is converted to reddish betacyanins such as betanin through a *GT*-catalysed reaction with glucose ^6,7^.

**FIGURE 1.**
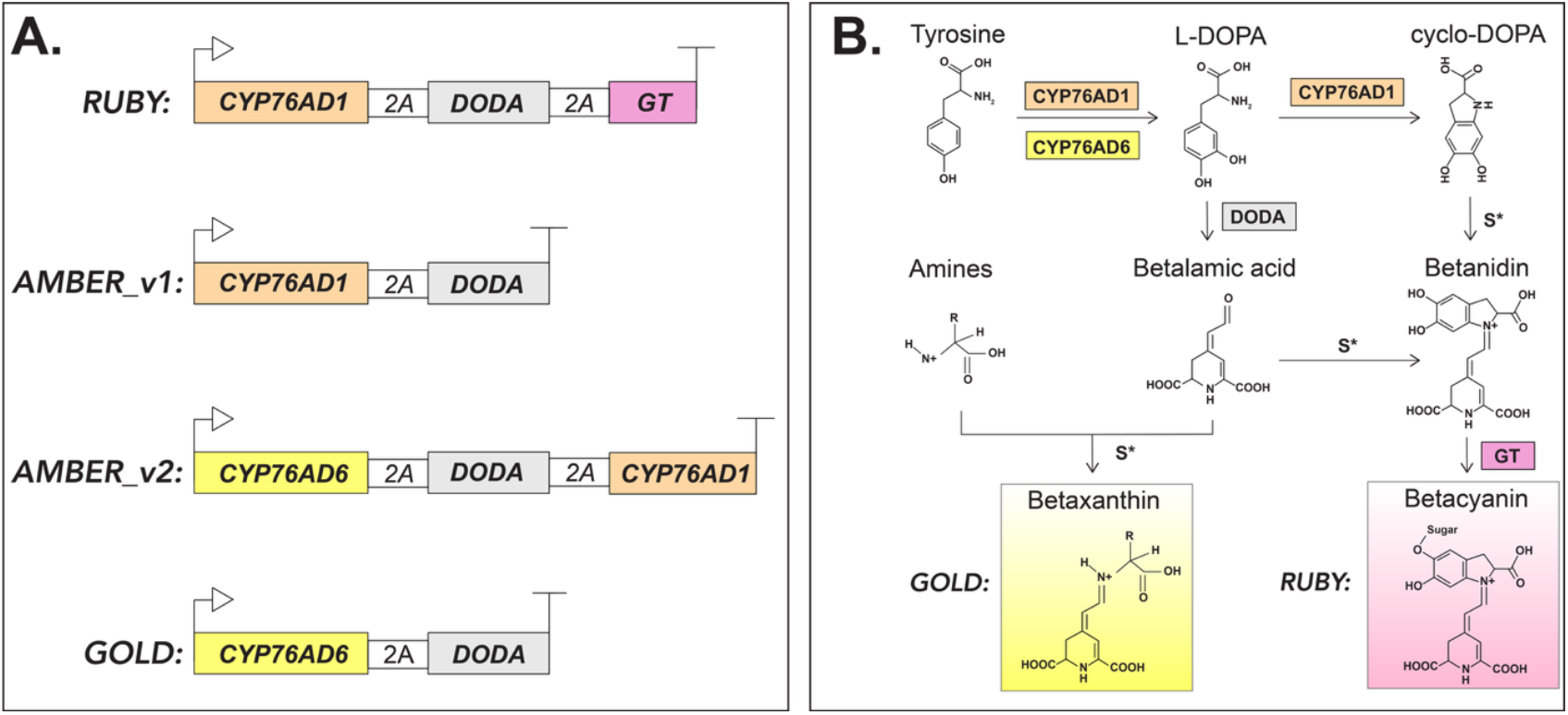
Genetic constructs and betalain biosynthesis pathway. (A.) Schematics of genetic constructs of *RUBY, AMBER_v1, AMBER_v2*, and *GOLD*. All four constructs contain combinations of *CYP76AD1, CYP76AD6, DODA*, and *GT* in open reading frames separated by 2A peptides that undergo self-cleavage, releasing the individual enzymes for betalain synthesis. (B.) Betalain biosynthesis pathway showing the role of the enzymes encoded in the four constructs. Whereas *RUBY* contains *GT* to generate betacyanin, the *AMBER* versions and *GOLD* generate predominantly betaxanthins from spontaneous reactions (S*) between amines and betalamic acid.

Thanks to its polycistronic single-gene architecture, RUBY has proven practical for molecular cloning and has been widely used to develop reporter systems for diverse applications. It has also served as a valuable tool for teaching complex biological and biotechnological concepts ^8,9^. Given its beetroot-red pigmentation, RUBY has also inspired artistic creativity, for instance enabling striking floral patterns facilitating their aesthetic design ^10,11^. However, to date, the “palette” of polycistronic genes encoding additional colors has been limited.

Several studies have demonstrated that other genes in the betalain pathway, particularly *CYP76AD6*, can also be used to produce yellow betaxanthins ^4,5,12–17^. Unlike *CYP76AD1, CYP76AD6* does not oxidize L-DOPA to cyclo-DOPA, and therefore most L-DOPA is converted to betalamic acid which can spontaneously undergo condensation reactions with amines and amino acids to form yellow, fluorescent betaxanthins. Here, we develop three polycistronic genes, *AMBER_v1, AMBER_v2*, and *GOLD* with various combinations of *CYP76AD1*/*CYP76AD6* and *DODA* separated by the self-cleavable 2A linkers to produce yellow, fluorescent betaxanthins in plant tissues.

## RESULTS

To produce yellow, fluorescent betaxanthin reporters and enable complex floral designs, we sought to express the biosynthetic pathway under a single promoter. To this end, we cloned three constructs as Golden Gate level 0 coding sequence parts, *AMBER_v1, AMBER_v2*, and *GOLD* containing either *CYP76AD1-2A-DODA, CYP76AD6-2A-DODA-2A-CYP76AD1*, or *CYP76AD6-2A-DODA* respectively (Figure 1). To test their ability to produce betaxanthins, we fused the coding sequences to a 35S promoter and terminator and transiently expressed them in *Nicotiana benthamiana* leaves using agroinfiltration ^18^. All three constructs produced visible orange-brown or yellow pigmentation within three days of infiltration that could be easily seen when leaves were cleared in ethanol (Figure S1).

We also took advantage of the petunia petal agroinfiltration assay to further evaluate the new reporters and quantify their colors ^19^. To achieve this, we transiently expressed the constructs described above in white petunia petals and observed visible pigmentation within 48 hours (Movie S1). *RUBY* produced a light purplish pink, AMBER_v1 and AMBER_v2 varied in color, with the average being a pale orangish yellow, and *GOLD* consistently produced yellow. Four days after infiltration, petal color was quantified with a colorimeter to achieve CIELAB values for each construct (Figure 2A). Because betaxanthins are fluorescent, we imaged the flowers using 460nm excitation light and a 520 ± 20 nm emission filter. Whereas *RUBY* fluorescence intensity was similar to non-infiltrated petals, *AMBER_v1* and *AMBER_v2* petals had on average approximately 10-fold higher fluorescence intensity than *RUBY* petals (n=10). Furthermore, *GOLD* produced on average >20 times stronger fluorescence intensity than *RUBY*.

**Figure 2.**
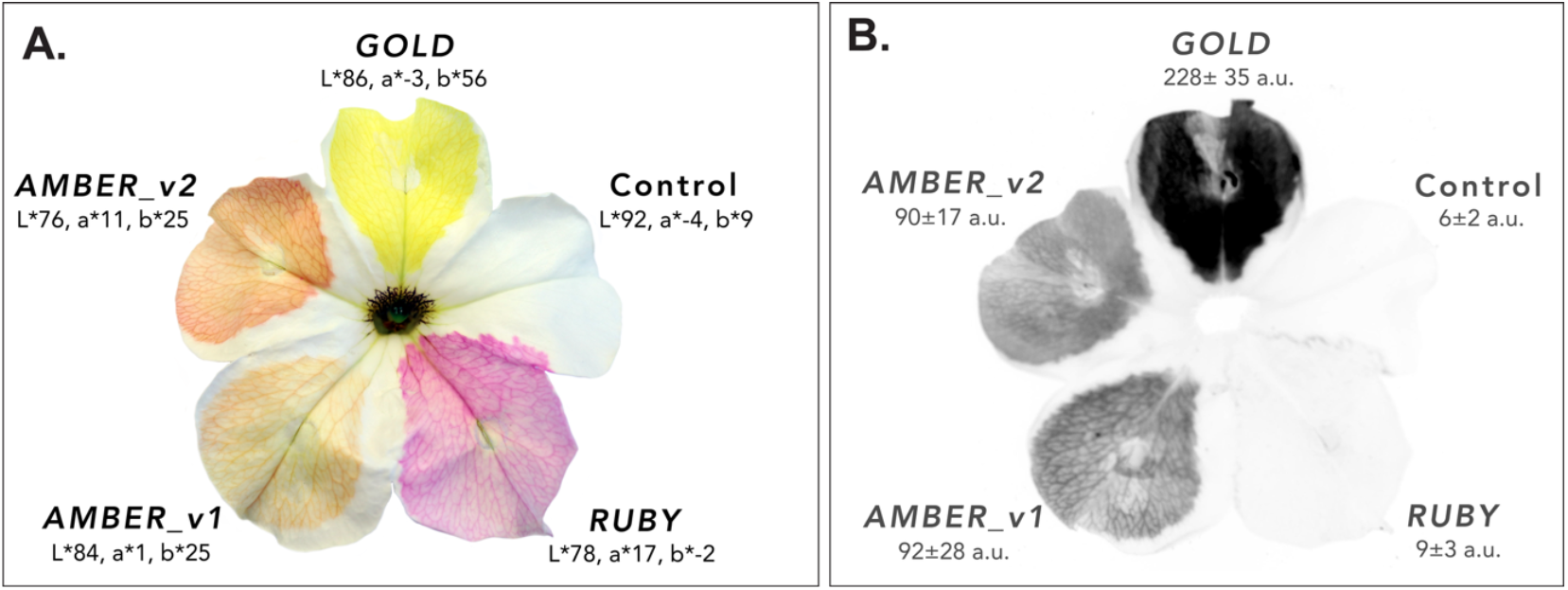
Colorimetric and fluorescent properties of constructs in *Petunia* petals. (A.) Petunia flower 4 days after infiltration of each petal with either *RUBY, AMBER_v1, AMBER_v2, GOLD*, or none (control) under a 35S promoter. Petal color from 5 independent flower infiltrations was analysed with a colorimeter showing the following CIELAB values: *RUBY* (l*= 78 ± 7, a*= 17 ± 6, b*= −2 ± 3), *AMBER_v1* (l*= 84 ± 2, a*= 1 ± 2, b*= 25 ± 7), *AMBER_v2* (l*= 76 ± 1, a*= 11 ± 2, b*= 25 ± 5), *GOLD* (l*= 85 ± 1, a*= −3 ± 2, b*= 56 ± 9), Control (l*= 92 ± 3, a*= −4 ± 1, b*= 9 ± 2).

To confirm that the yellow pigmentation and fluorescent properties of *AMBER* and *GOLD* were due to the production of betaxanthins, we performed high-performance liquid chromatography-mass spectrometry (HPLC-MS) on water-soluble extracts from ethanol-cleared *N. benthamiana* leaves infiltrated with the four constructs. Whereas the *RUBY* extract showed a very prominent chromatogram peak with the expected spectral properties and mass of betanin, *AMBER_v1, AMBER_v2*, and *GOLD* had little or no betanin, but other peaks (Figure 3A). These other compounds had UV/visible spectra and masses like those previously described for betaxanthins ^4,5^. Two prominent betaxanthins, Vulgaxanthin and Indicaxanthin were present in all four construct samples, with a greater abundance in *AMBER* samples compared to *RUBY* and even greater in the *GOLD* sample (Figure 3B,C).

**Figure 3.**
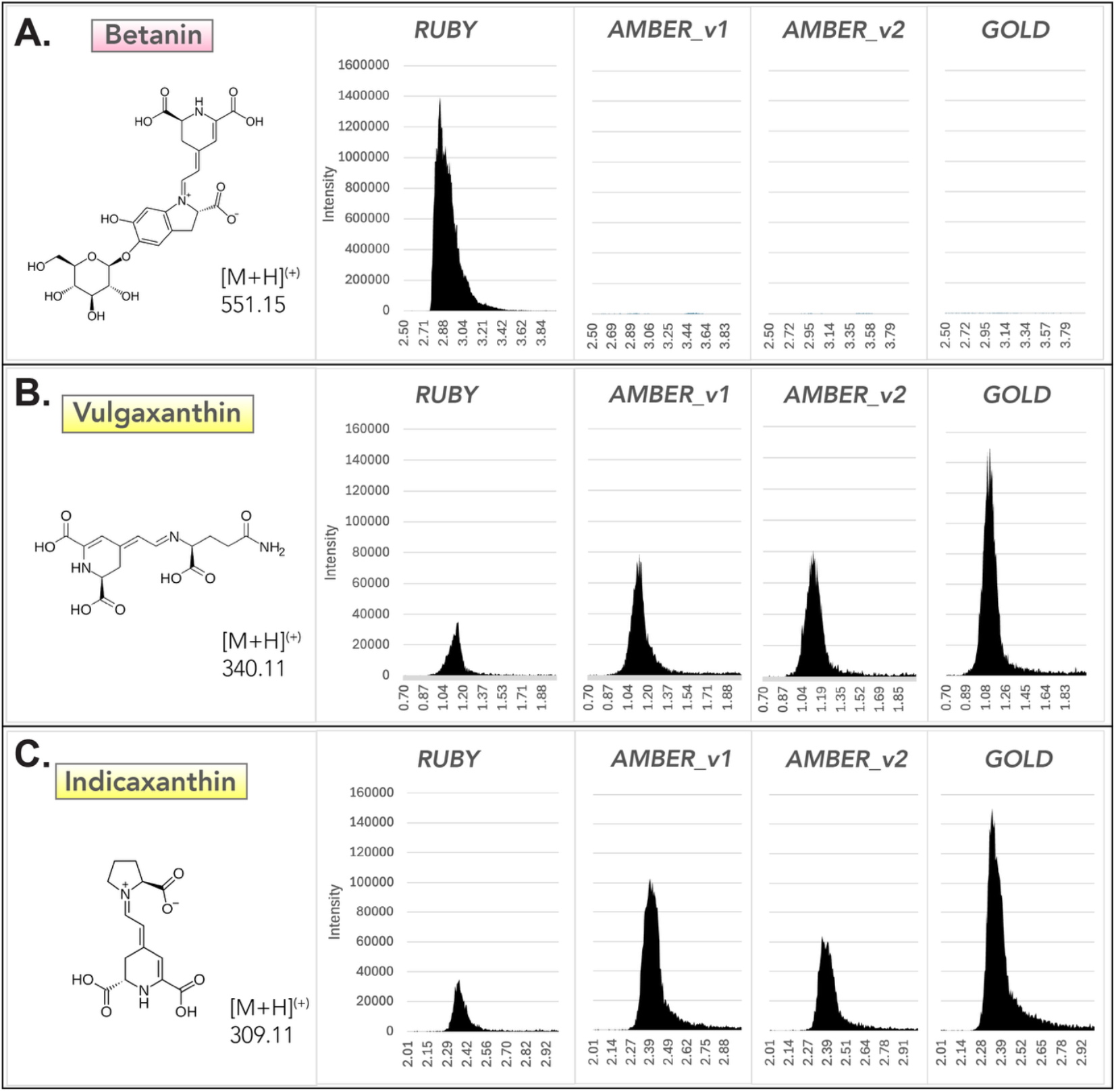
HPLC-MS of Betalain compounds from each construct. (A-C.) Extracted ion chromatograms of mass 551.15 (expected [M+H]^(+)^ for Betanin), mass 340.11 (expected [M+H]^(+)^ for Vulgaxanthin), and mass 309.11 (expected [M+H]^(+)^ for Indicaxanthin) from leaf disks expressing each construct. Betanin was only prevalent in leaves expressing *RUBY*, whereas Vulgaxanthin and Indicaxanthin were most prevalent in leaves expressing *GOLD*, followed by *AMBER_v1* and *AMBER_v2*, and *RUBY*. Y axis shows signal intensity and X axis retention time in minutes. All graphs have the same Y-axis and X-axis scales.

## DISCUSSION

Color is among the most accessible and interpretable outputs of biological systems as it can be directly observed without specialized instruments. As such, visual color markers have broad utility in research, education, and art. Historically, color phenotypes have played a pivotal role in foundational biological discoveries, including the identification of transposable elements ^20^, sex-linked inheritance ^21^, and RNA interference ^22^.

Beyond their function as gene expression reporters, genetically encoded pigments offer unique opportunities for artistic and educational applications. Bacterial chromoproteins, for example, have been used to create living “paintings” and serve as engaging tools for science outreach and teaching ^23^.

Flowers, unlike bacteria, have a long history as subjects of art and as ornamental objects, presenting a compelling canvas for genetic design aimed at manipulating pigmentation. Here, we expand the toolkit of polycistronic genes for genetically encoded pigments in flowers to now include *AMBER* and *GOLD*. We envision their use in diverse contexts spanning research, education, and creative expression.

## MATERIAL AVAILABILITY

The plasmids described in this study can be provided upon request to the corresponding author (ND). Plasmids are being submitted to Addgene.

## MATERIALS & METHODS

### Construct cloning

*RUBY, AMBER_v1, AMBER_v2*, and *GOLD* were cloned as level 0 CDS golden-gate parts using the PCRs and synthetic fragments described in table 1. PCRs were performed using KAPA HiFi Hotstart ready mix and gel extractions performed using Zymoclean columns with 2 x 6ul elutions. Equimolar ratios of fragments were shuttled into the acceptor pICSL41308_CP (AATG – GCTT overhangs) at a 2:1 molar ratio to the acceptor, using standard golden-gate cycling protocols with 3 min digestions with BpiI (37°C), 4 min ligations (16°C) x 26 cycles.

*35S:RUBY, 35S:AMBER_v1, 35S:AMBER_v2*, and *35S:GOLD* were assembled as level 1 golden-gate transcriptional units, using the level 1 acceptor pICH47742_RFP, short *35S* promoter pICH51277, and *35S* terminator pICH41414.

### Transientexpression assays

Plasmids were transformed into *Agrobacterium tumefaciens* strain GV3101 with electroporation and grown overnight as 5ml cultures with appropriate antibiotics at 28°C until saturation. Cells were harvested by centrifugation at 4000 x g at room temperature for 10 minutes before being resuspended in infiltration buffer (10 mM MgCl2, 10 mM MES-KOH pH 5.6, 200 µM acetosyringone) to OD_600_ 0.25. Suspensions were co-infiltrated with the Tomato bushy stunt virus (TBSV) silencing inhibitor p19 into either 4-5 week old *Nicotiana benthamiana* leaves or Mitchel line W115 petunia petals of newly opened flowers ^24,25,19^. Petunia flowers were cut at the pedicle before infiltration and left in 50ml falcon tubes with water for the duration of incubation.

### Photography and Timelapse imaging

Photo in Figure 2A was taken with a Canon EOS RP and RF 35mm f/1.8 STM lens with white paper as a background. “Auto-tone” function was applied and background manually removed in photoshop. Adjustments were applied uniformly to the whole image.

The timelapse in Movie S1 was taken with a Nikon D3200 (AF-S 18-105mm objective) whose shutter pin was connected to an Arduino board programmed to turn the lights on in the growth chamber for 5 seconds and take a picture every 5 minutes. This enables consistent lighting throughout the photoperiod of 16-hour daylight 8-hour night. The images were then assembled in Adobe Premiere Pro followed by time and lighting adjustments in Adobe After Effects.

### Colorimetry and Fluorescence analysis

For testing fluorescence of petunia petal infiltrations, 10 flowers were imaged 4 days post infiltration using an Amersham ImageQuant 800 on fluorescence mode using 460nm excitation light and 520± 20 nm Cy2 emission filter. Images were converted to 8-bit in Fiji and a 100px roi circle measured for signal intensity of each petal.

For colorimetry 5 independently infiltrated petals per construct were cut out and measured on a HunterLab UltraScan VIS colorimeter in reflectance mode with specular excluded.

### Analytical chemistry

Betalains were extracted from *Nicotiana benthamiana* leaves 4 days post-infiltrated with each construct. Leaves were first cleared in ethanol overnight and then 500mg flash frozen in liquid nitrogen before grinding in 5ml H_2_O with mortar and pestle. Samples were centrifuged and supernatant transferred to tubes for HPLC-MS analysis.

Samples were run on an Agilent Infinity II UHPLC system with a G6546A Q-ToF mass spectrometer. Separation was on a 100×2.1mm 2.6µ Kinetex EVO C18 column (Phenomenex) using a gradient of 1% - 95% acetonitrile versus 0.1% formic acid in water, run at 40°C and 600µL.min^−1^. MS data was collected in both positive and negative mode in separate runs. The mass spec collected full spectra using positive mode electrospray ionisation from *m*/*z* 100-1700 at 125msec per spectrum, and also auto MS/MS (data-dependent MS2) of the two most abundant precursor ions, also at 125msec per spectrum (or 25000 counts, whichever came first), with medium (*m*/*z* 4.0) isolation width and 35% collision energy. After a precursor had been selected twice, it was excluded for 0.1min in favour of less abundant precursors. The system was set to recognise isotope peaks using the common organic molecules model. Spray chamber conditions for the JetStream source were 320°C drying gas at 8L.min^−1^, 35psi nebulizer pressure, 350°C sheath gas at 11L.min^−1^, 120V fragmentor, 3500V Vcap, 1000V nozzle voltage. The instrument used lock-masses at *m*/*z* 922.0098, 121.0509 in positive mode, and *m*/*z* 1033.9881, 112.9859 in negative mode.

Separation was on a 100×2.1mm 2.6µ Kinetex EVO C18 column (Phenomenex) using the following gradient of acetonitrile (solvent B) versus 0.1% formic acid in water (solvent A), run at 40°C and 600µL.min^−1^: 0 min, 1% B; 1 min, 1% B; 5.5 min, 30% B; 7.0 min, 95% B; 8.0 min, 12% B followed by re-equilibration.

## Supporting information

HPLC-MS, CIELAB, Fluorescence Quantification

## ACKNOWLEDGEMENTS

We thank Francesca M. Quattrocchio for providing the W115 Petunia line seeds. We thank all members of the TSL Support Services for their invaluable assistance and Maryam Sargolzaei for help with petal infiltrations.

## AUTHOR CONTRIBUTIONS

N.D. conceived and performed all experiments, prepared the figures, and wrote the paper. M.Y. cloned the level 0 Golden Gate plasmid parts. L.H. performed the HPLC-MS and data analysis. S.K. administered the project and helped write the paper. N.D. and S.K. acquired funding.

