## Supplementary Figures, Table, and Movie Legend for "*AMBER* and *GOLD*: Polycistronic Genes for Betaxanthin Production in Plants"

### Supplementary Information

[Figure S1, Table S1, Movie S1 legend]

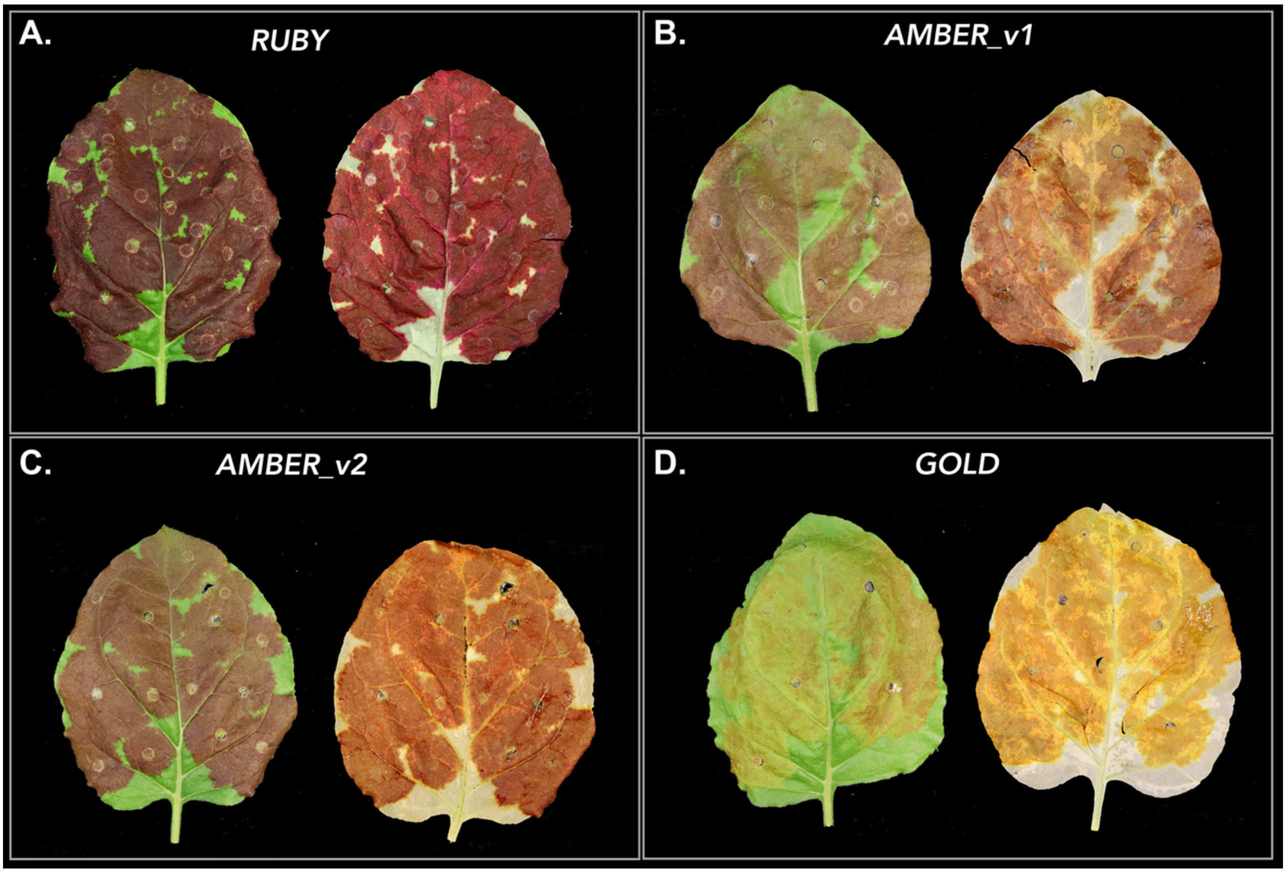


**Figure S1. Leaves infiltrated with each construct**

(A-D.) *Nicotiana benthamiana* leaves four days post-infiltration with each construct before clearing (left) and after ethanol clearing (right).

| **Table 1.** Primers, PCRs, and Synthetic Fragments used in this study |
|---|
| *RUBY* (pICSL80094)<br>PCR: 3962bp<br>Template: Addgene (#160908)<br>Primers:<br>MY_RUBY_AATG_S:<br>GGAAGACTTAATGGATCATGCGACCCTCGCCATGATCCTCGCG<br>MY_RUBY_GCTT_AS:<br>GGAAGACTTAAGCTCACTATCACTGGAGGCTTGGCTCAAGTTTGG |
| *AMBER_v1* (pICSL80187)<br>PCR: 2390bp<br>Template: pICSL80094<br>Primers:<br>MY_RUBY_AATG_S:<br>GGAAGACTTAATGGATCATGCGACCCTCGCCATGATCCTCGCG<br>MY_ND_DODA1_GCTT_AS:<br>GGAAGACTTAAGCTTAGGCGGAGGTGAACTTGTAGGAGCCGTGGC |
| *AMBER_v2* (pICSL80189)<br>Synthetic Fragment + PCR1 + PCR2<br><br>PCR1: 940bp<br>Template: pICSL80094<br>Primers:<br>MY_ND_P2A1_TAGC_S:<br>GGAAGACTTTAGCGGAGCTACCAATTTTAGCCTCCTTAAGCAG<br>MY_ND_P2A2_CGAG_AS:<br>GGAAGACTTCTCGACATCTCCTGCTTGCTTGAGCAGGCTAAAG<br><br>PCR2: 1517bp<br>Template: pICSL80094<br>Primers:<br>MY_ND_CYP-AD1_CGAG_S:<br>GGAAGACTTCGAGGAAAATCCTGGCCCCATGGATCATGCGACCCTCGCCATGATCCTCGC<br>MY_ND_CYP-AD1_GCTT_AS<br>GGAAGACTTAAGCTTAGTAGCGCGGAATCGGGATGAGCTTGAGCGGC |
| *GOLD* (pICSL80188)<br>Synthetic Fragment + PCR<br><br>PCR: 896bp |

Template: pICSL80094
Primers: MY_ND_P2A1_TAGC_S:
GGAAGACTTTAGCGGAGCTACCAATTTTAGCCTCCTTAAGCAG
MY_ND_DODA1_GCTT_AS:
GGAAGACTTAAGCTTAGGCGGAGGTGAACTTGTAGGAGCCGTGGC

Synthetic Fragment (*Beta vulgaris CYP76AD6_2A*)

GGAAGACTTAATGGATAACGCAACACTTGCTGTGATCCTTTCCATTTTGTTTGTGTTTTACCACATTTTCA
AATCCTTTTTCACCAATTCTTCATCTCGTAGGCTTCCTCCTGGTCCCAAACCCGTGCCAATTTTTGGCAAC
ATTTTCGATCTTGGCGAAAAGCCTCATCGATCTTTTGCCAATCTATCTAAAATTCACGGCCCTTTGATTAG
CCTAAAGTTAGGAAGTGTAACAACTATTGTTGTTTCCTCGGCCTCTGTGGCCGAGGAAATGTTCCTTAAAA
ATGACCAAGCACTTGCTAACCGAACCATTCCTGACTCGGTTAGGGCTGGTGACCACGACAAATTATCCATG
TCGTGGTTGCCTGTTTCCCAAAAATGGAGAAATATGAGAAAAATCTCCGCTGTCCAATTACTCTCCAACCA
AAAACTTGATGCTAGTCAACCTCTTAGACAAGCTAAGGTGAAACAACTTTTATCATACGTACAAGTTTGTT
CCGAAAAAATGCAACCCGTCGATATTGGACGGGCCGCATTTACAACGTCACTTAATTTATTATCAAACACA
TTTTTCTCAATCGAATTAGCAAGTCATGAATCTAGTGCTTCCCAAGAGTTTAAACAACTCATGTGGAATAT
TATGGAGGAAATTGGAAGGCCTAATTATGCTGATTTTTTCCCTATTCTTGGTTACATTGATCCCTTTGGTA
TAAGACGTCGTTTGGCTGGTTACTTTGATAAACTCATTGATGTTTTCCAAGACATTATTCGTGAAAGACAA
AAGCTTCGATCTTCTAATTCTTCCGGCGCAAAACAAACAAATGACATTCTTGATACTCTTCTTAAACTCCA
TGAAGATAATGAGTTGAGTATGCCTGAAATTAATCACCTTCTCGTGGATATCTTTGACGCCGGAACAGACA
CAACAGCAAGCACATTAGAATGGGCGATGGCCGAACTTGTGAAAAACCCGGAAATGATGACTAAAGTTCAA
ATTGAAATCGAACAAGCTCTTGGAAAAGATTGCTTAGACATACAAGAATCCGACATCTCAAAACTACCTTA
TTTACAAGCCATTATAAAAGAAACGTTACGTTTACACCCTCCTACTGTGTTTTTGCTGCCTCGAAAGGCAG
ACAATGACGTAGAGTTATATGGCTACGTTGTACCAAAGAATGCTCAAGTCCTTGTCAATCTTTGGGCAATT
GGTCGTGATCCAAAGGTATGGAAAAATCCGGAAGTATTTTCTCCTGAAAGGTTTTTAGATTGCAATATCGA
TTATAAAGGACGAGATTTCGAACTTTTACCCTTTGGTGCTGGTAGAAGGATATGCCCTGGACTTACTTTGG
CATATAGAATGTTGAACTTGATGTTGGCTACTCTTCTTCAAAACTACAATTGGAAACTTGAAGATGGTATC
AATCCTAAGGATTTAGACATGGATGAGAAATTTGGGATTACATTGCAAAAGGTTAAACCTCTTCAAGTTAT
TCCAGTTCCCAGAAACGGTAGCGGAGCTACTTGTCTTCC

**Movie S1.** Timelapse of petunia petal infiltration with constructs.

Looped 24-hour timelapse of petunia flowers infiltrated with *RUBY* (top petal), *GOLD* (bottom left petal), and *AMBER_v2* (bottom right petal) one day after infiltration.
